# Cortical and Cardiovascular Regulation Support Adaptation to Ambient Sound Environment

**DOI:** 10.1101/513937

**Authors:** Yu Hao, Disha Gupta, Qiuyan Sun, Lin Yao

**Affiliations:** Department of Design and Environmental Analysis, Cornell University.; School of Medicine, New York University.; Department of Nutritional Science, Cornell University.; School of Electrical and Computer Engineering, Cornell University.

**Author notes:** Correspondence to: Yu Hao,.

**Keywords:** Electroencephalograph, mismatch negativity, biological regulation, unattended sound, adaptation, attention, heart-rate variability

## Abstract

Adaptation in complex biological systems is a consequence of sensitivity to environmental changes. We investigate how responses to unattended background sound changes are influenced by the interaction of external (pitch sequence context) and internal factors (biological regulation). Cortical responses (electroencephalograph Mismatch Negativity) demonstrated increased intensity and prolonged latency in unattended pitch changes in the context of the ascending pitch sequence compared with the descending pitch sequence. Moreover, ascending pitch context is associated with more activation of anterior cingula cortex and insula, suggesting arousal effect and internal regulation. However, the intensified and prolonged responses were supported by cortical and cardiovascular biomarkers for regulation respectively. These findings suggest that biological regulation may be associated with environmental context and play different roles in response sensitivity in terms of intensity and speed.

---

With the dynamic nature of the ambient environment, organisms as ensembles of both brain and body interact with the environment while maintaining equilibrium through adaptation and regulation ^1,2^. Although it is generally accepted that this adaptation results from the integration of brain, body and environment ^2^, the neural basis in this system is less well understood. One role that the brain plays is the modulation of perceptual sensitivity based on contextual information from the environment. For example, when stimuli are processed attentively, perception of a tone pitch was biased towards previously heard pitches, reflecting the temporal binding of successive frequency components ^3^. When stimuli are processed unattentively, regular context was associated with higher neural response to changes than random context, indicating that the brain is more sensitive to change in a predictable context than in a random context ^4^. This responsiveness to environmental changes is essential for survival, especially when changes are not always in the focus of attention. Awareness of surroundings is enabled by perceptual processes which bring into play autonomic and neuroendocrine systems that enable responses to emergencies ^2^. However, it remains unclear how the brain responds to changes in *unattended* stimuli in different dynamic environmental contexts and in what way the brain and body work together to regulate response sensitivity to such stimuli. Sensitivity to stimuli has usually been shown to be reflected in a quicker and stronger neural response. Thus, the purpose of this study is to examine how speed and intensity of detecting unattended changes are regulated and how they are associated with different contexts, applying one neural regulation biomarker from the brain and one somatic regulation biomarker from the heart.

In this article, we study unattended sensory information processing by measuring neural responses to ambient acoustic stimuli, following the well-established experiment paradigm of electroencephalograph (EEG) based Mismatch Negativity (MMN). MMN is an electrophysiological response that is evoked by an oddball or deviant event in a sequence of repeated or familiar events (the standards), which might reflect the borderline between automatic and attention-dependent processes ^5^. The MMN is generally observed in the temporal and frontal areas at peaks around 100-250ms from change onset ^6^. When immersed in ambient environmental stimuli, the brain actively predicts sensory inputs and compares the incoming stimuli with these top-down predictions, eliciting an MMN when a prediction error is detected ^6,7^. However, most studies have used static parameters and it remains unclear how stimuli with rapidly changing context is processed. To test such a contextual effect, twenty participants were individually immersed in auditory odd-ball streams that were either rapidly ascending in pitch (600-1400 Hz) or descending in pitch (1400Hz-600 Hz). The odd ball in each stream was the last pitch, 1600 instead of 1400 Hz in an ascending stream and 400 Hz instead of 600 Hz in a descending stream (Fig. 1a). To distract attention from the auditory stimuli, the participants were asked to watch a self-selected silent documentary. The primary aim was to assess the effect of context on the MMN response in terms of intensity (i.e., amplitude) and speed (i.e., latency, the timing from stimulus onset to response peak).

**Figure 1.**
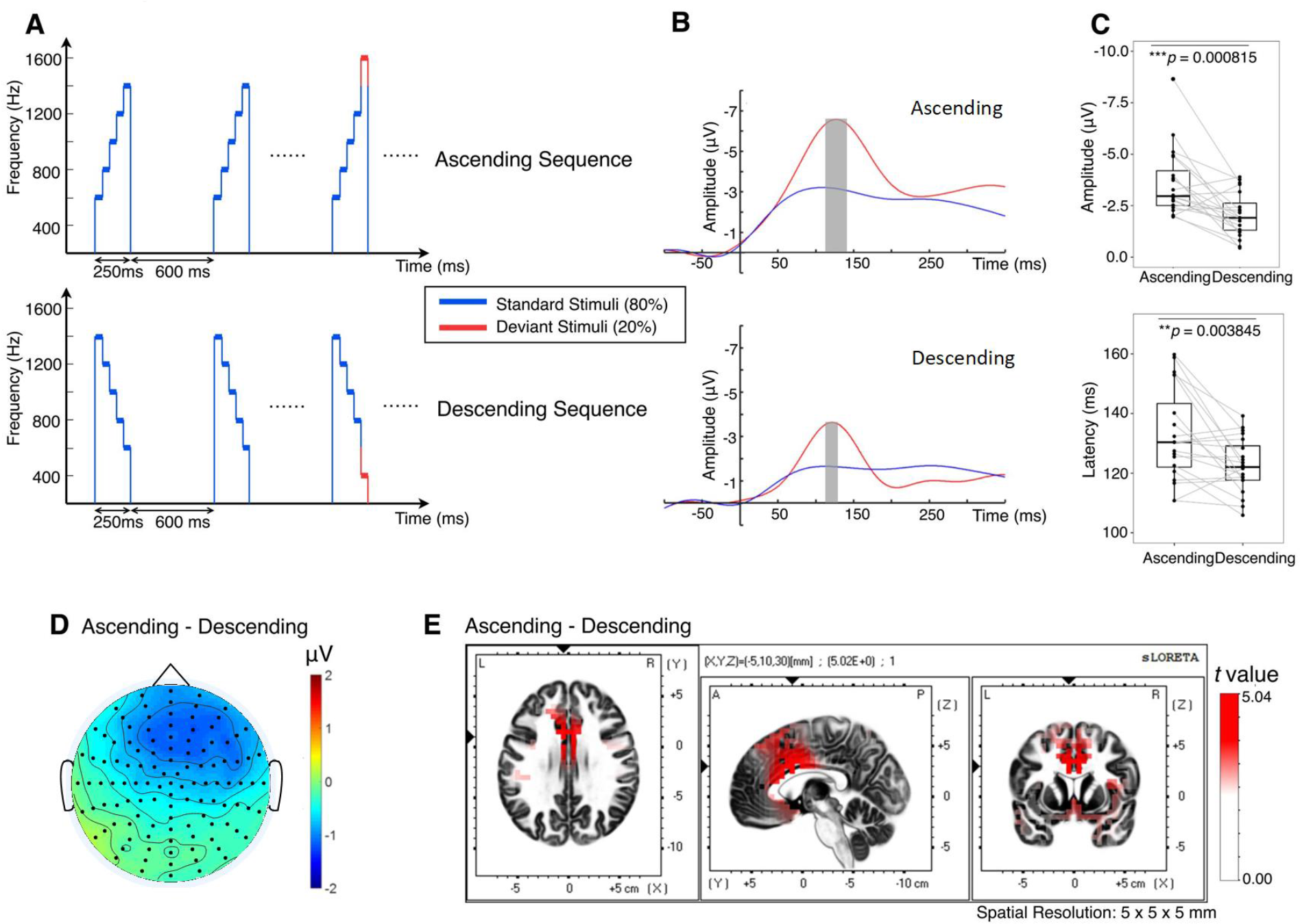
Contextual effect of unattended sound changes on brain response. (**a**) Experimental paradigm. The time interval between two stimuli was 600 ms and each stimulus (250 ms long) contained 5 concatenated tones (50 ms each) with either ascending or descending pitch. Twenty percent of the stimuli set in either condition had a deviant tone sequence (plotted in red)— a 200 Hz additional pitch increment or decrement tone in substitution to the fifth tone of the standard stimuli, in which pitch increment or decrement between each tone was constantly 200 Hz. (**b)** Grand average MMN at electrodes having largest difference between standard and deviant in the fronto-central region in two pitch sequence conditions. Grey bars are peak intervals (peak time ± SD) for ascending sequence at 118-149 ms and descending sequence at 114-132 ms. (**c)** Ascending sequence elicited greater MMN amplitude (−3.60 ± 1.65 *μV*) than descending sequence (−2.08 ± 1.06 *μV*). The MMN latency was greater in the ascending sequence (132.84 ± 15.53 ms) than in the descending sequence (122.55 ± 9.15 ms). (**d**) Scalp topography of MMN difference between ascending and descending condition in their respective peak intervals, which exhibits in the fronto-central region. (**e**) Significant increase in current source density of the MMNs in the ascending condition compared with the descending condition at the respective peak intervals at regions such as superior temporal gyrus, inferior frontal gyrus, medial frontal gyrus, anterior cingulate, cingulate gyrus and insula. *t* > 2.093 indicates significant differences in paired *t*-test by whole scalp multiple comparison correction.

Prior work indicates that the pitch of environmental sounds alters perceived valence and arousal of auditory stimuli ^8–10^. The stimuli were designed to have equal intensity, but the higher pitch stimuli would also be perceived as louder and the descending as softer, as per loudness-frequency curves. The pitch and loudness are fundamental characteristics of speech that embody emotional context ^11^. The regular pattern of the ambient sound might generate a baseline environmentally induced affective state. Thus, we hypothesized that arousal and biological regulation mediate contextual effect on response to unattended changes. Usually, MMN latency and amplitude have been found to be coupled, with a quicker (smaller latency) and a larger (greater amplitude) MMN response indicative of sensitivity to change, both because of variations in deviant stimuli and cognitive decline ^12–15^. Therefore, the second aim was to understand MMN response intensity and speed in relation to arousal and biological regulation processes. Biological regulation involves volitional and nonvolitional components enabled by executive functioning and the autonomous system ^22,23^.

## Results

We first analyzed the MMN elicited by the oddball/deviant pitch, obtained by subtracting the response to the standard stimulus (blue) from the response to the deviant stimulus (red) (Fig. 1b). The ascending sequence elicited a slower (*t* (19) = 3.29, *p* = 0.003845, Cohen’s *d*_*z*_ = 0.74, Power (*1- β*) = 0.88) but larger (*t* (19) = −3.97, *p* = 0.000815, Cohen’s *d*_*z*_ = 0.89, Power (*1- β*) = 0.96) MMN than the descending sequence (Fig. 1c). This phenomenon is different from previous findings wherein the latency and the amplitude of MMN were coupled that together reflect the sensitivity to change ^12–15^. If deviants in ascending condition were simply more salient than in descending, they would be expected to elicit greater amplitude and smaller (or similar) latency. Thus, these data suggest 1) detection intensity (MMN amplitude) and speed may reflect different aspects of sensitivity, which were possibly processed by different mechanisms, and 2) when exposed to different environmental contexts, these mechanisms could be differentiated. Moreover, the MMN latency and amplitude did not correlate with each other in any condition or cross conditions (Fig. s1). This finding provides further support that deviant stimuli with higher pitch not only influence sensitivity but are also associated with the interaction of internal regulation with external context. We therefore sought to discriminate between these two external contexts by brain activations.

We next analyzed and compared the current source density of the MMNs between the ascending condition and the descending condition. Superior temporal gyrus and inferior frontal gyrus were significantly higher in MMN elicited during ascending condition than in the descending condition, *p* <.01 (Fig. 1d). This may indicate that deviants in the ascending condition were more salient than deviants in the descending condition, which suggests a bias in top-down processing of attention towards novelty ^16^. Current source density of MMN in the ascending condition was also significantly higher than descending condition in the insula, *p* <.01 and in the cingulate gyrus including anterior cingulate cortex (ACC), *p* <.001 (Fig. 1d). These results are consistent with the error detection mechanism mentioned above as the anterior cingulate cortex contributes to attention, which serves to regulate both cognitive and emotional processing ^17,18^. On the other hand, the insula receives viscerosensory inputs and responsible for affective experience, particularly in negative emotion suppression ^19–21^. EEG source localization results should be interpreted with caution, however, because its spatial resolution is not equivalent to that of functional MRI. Altogether, the results indicate that the brain is more sensitive to changes in ascending sequence and executed more regulation procedures, but do not explain the discrepancy results of MMN amplitude and latency.

We then examined how MMN amplitude and latency were modulated by regulation biomarkers under both contexts. Biological regulation involves volitional and nonvolitional components enabled by executive functioning and the autonomous system ^22,23^. As a neural biomarker of executive functioning, EEG frontal theta/beta ratio is an electrophysiological marker for prefrontal cortex-mediated executive control over attentional and emotional information, with lower frontal theta/beta ratio representing better focused attention and emotion regulation ^24,25^. As a somatic marker of autonomic regulation, heart-rate variability (HRV) reflects stress and regulated emotional response, with higher values representing better regulation ^26^. We estimated HRV with the root mean square of the successive differences (RMSSD) score.

We found that both theta/beta ratio (6.001 ± 2.405) and RMSSD (0.049 ± 0.011s) were not significantly different between ascending and descending sequence conditions (Fig. s2). However, a strong nonlinear correlation existed between theta/beta ratio and RMSSD (*Spearman* nonlinear correlation test: *ρ* = 0.4259, *S* = 6120, *p* = 0.006549): moderate theta/beta ratio corresponded to the highest HRV and, as theta/beta ratio increased, HRV declined (Fig. 2a). This suggests that frontal theta/beta ratio and HRV may represent different aspects of regulation. The data showed that in the ascending sequence condition, higher theta/beta ratio was associated with smaller MMN latency and higher HRV was associated with larger MMN amplitude, whereas no correlations were found in the descending sequence condition (Fig. 2 b-e). Moreover, higher HRV was also associated with smaller MMN latency across conditions, *χ*^2^(1) = 3.70, *p* = 0.05451 (Fig. 2d). Statistical interaction modeling showed that theta/beta ratio interacted with condition on MMN latency, *χ*^2^(1) = 5.10, *p* = 0.0238958, *PCV* (proportion change of variance) = 14.92% (Fig. 2b). HRV did not interact with condition on MMN latency, *χ*^2^(1) = 0.24, *p* = 0.6234, *PCV* = −7.72% (Fig. 2c). HRV marginally interacted with condition on MMN amplitude, *χ*^2^(1) = 3.67, *p* = 0.05548, *PCV* = 10.19% (Fig. 2d). Theta/beta ratio did not interact with condition on MMN amplitude, *χ*^2^(1) = 0.04, *p* = 0.8448, *PCV* = −2.06% (Fig. 2e). Taken together, these results show that biological regulation biomarkers can account for neural responses to unattended changes, but only under some specific conditions, such as the ascending pitch sequence context described in this study. Further, the response intensity and speed cannot be accounted for by the same biomarkers; rather they can be accounted for by cortical and cardiovascular biomarkers of regulation respectively.

**Figure 2.**
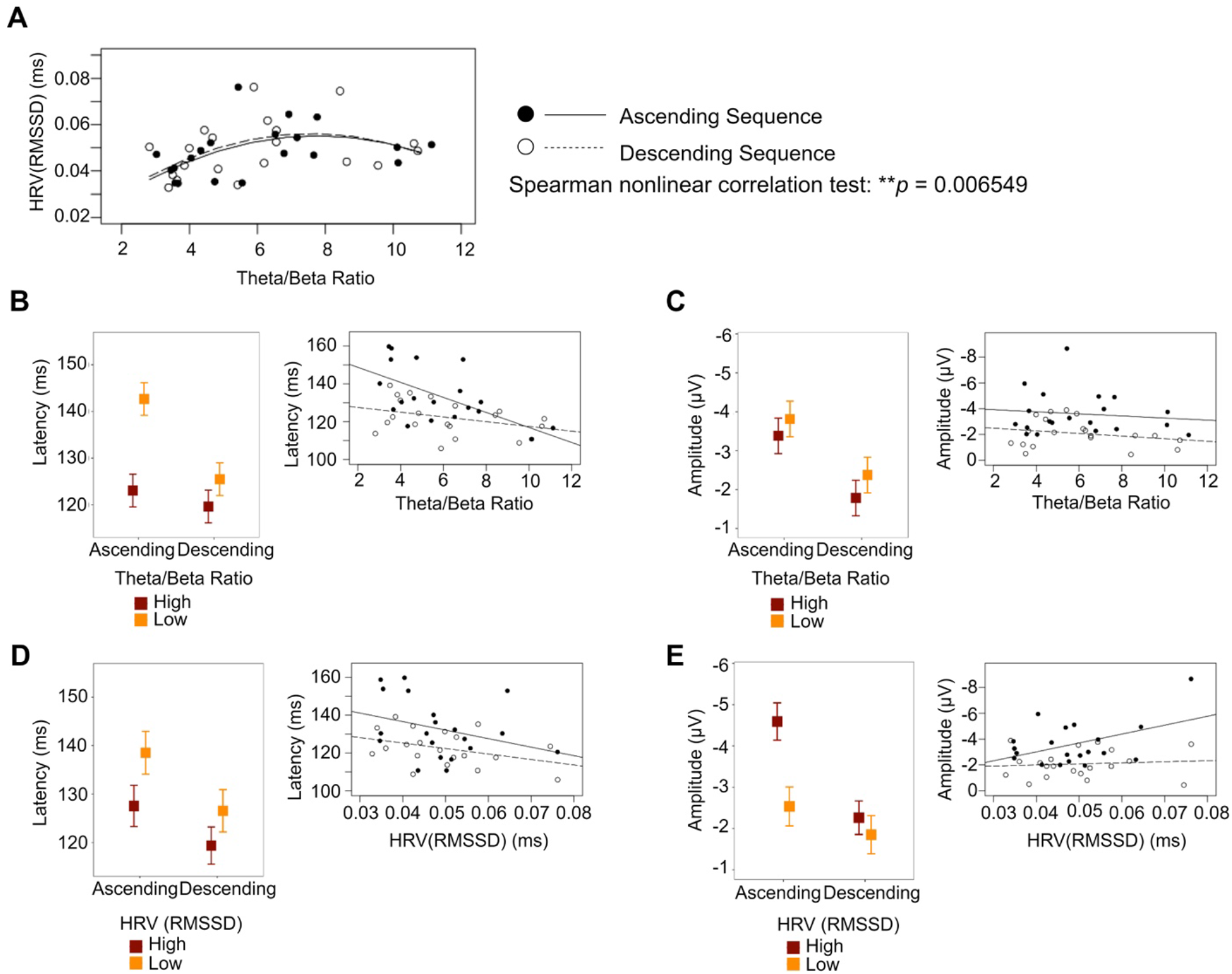
Contextual effect modulated by regulation biomarkers. Mixed effect models were applied to test the moderation effect of regulation biomarkers with experimental condition. High and low theta/beta ratio and HRV are plotted ± SD from the mean for descriptive purposes only in the Figures. Inferential analyses maintained the continuous nature of the variables. (**a**) A strong non-linear correlation was found between theta/beta ratio and HRV. **(b)** Theta/beta ratio interacted with condition on MMN latency. In the ascending condition, lower theta/beta ratio was associated with greater MMN latency, *r* = −0.623, *t* (18) = −3.378, ** *p* = 0.003347; whereas in the descending condition, theta/beta ratio was not correlated to MMN latency, *r*= −0.334, *t* (18) = −1.51, *p* = 0.1498. (**c**) Theta/beta ratio was not associated with MMN amplitude in either condition. In the ascending condition, *r* = 0.1194, *t* (18) = −0.51, *p* = 0.6161; in the descending condition, *r* = 0.2317, *t* (18) = −1.01, *p* = 0.3256. (**d**) HRV did not interact with condition on MMN latency but had a marginal main effect. In the ascending condition, *r* = −0.3152, *t* (18) = −1.41, *p* = 0.1759; in the descending condition, *r* = −0.3789, *t* (18) = −1.74, *p* = 0.0995. (**e**) HRV interacted with condition on MMN amplitude. In the ascending condition, greater HRV was associated with greater MMN amplitude, *r* = −0.453, *t* (18) = 2.16, * *p* = 0.0446; whereas in the descending condition, HRV was not correlated to MMN amplitude, *r* = −0.09218, *t* (18) = 0.39, *p* = 0.6991.

## Discussion

We investigated the interaction between the brain, body, and environment that contribute to neurophysiological responses to unattended changes in ambient acoustic stimuli. We have reported neurophysiological responses to unattended sound changes in two sound contexts: the ascending pitch sequence and the descending pitch sequence. We also showed how the response sensitivity in terms of speed and intensity was predicted by biological regulation biomarkers from brain and heart differently in both contexts. Overall, unattended sensory perception is influenced by context and sensitivity to changes is possibly related to emotion driven biological regulation. Apart from external context, biological regulation plays a key role in adaptation and formulate equilibrium in human-environment relations. The anterior cingulate and insula are involved in the neural circuitry of the central autonomic network. This network is a part of an internal regulation system wherein the brain controls visceromotor and neuroendocrine processes, essential for adapting environmental demands ^27^. As during the ascending condition, more activation in ACC and insula was observed; the context-related heightened arousal triggered more regulation, so that participants could adapt to unattended changes ^22,23^. Participants with higher HRV elicited quicker and stronger MMN responses, suggesting that these individuals could better anticipate and regulate unexpected environmental changes.

In other studies, researchers directed participants on whether or not to focus attention on experimental stimuli in order to contrast participants’ responses ^28–31^, but did not take the stimuli context into consideration. In our study, all participants were instructed to focus their attention on documentary videos instead of the ambient acoustic stimuli. However, their neural responses to deviant stimuli differed because individual attentional control as one of the regulation processes was manifested under the ascending sequence but not the descending sequence context. In our data, slower response was observed in individuals with low theta/beta ratio. These individuals were superior in attentional control during focused attention tasks ^24,32^ and might allocate more regulation effort to the elevated arousal experienced during the ascending sound condition. The low theta/beta ratio individual would be less sensitive to unattended stimuli changes. However, participants who were inferior in attentional control might be more susceptible to distraction. Poor attentional control in attentive stimuli are associated with higher theta/beta ratio in individuals diagnosed with ADHD or anxiety disorders ^25,32^. In unattended situations, they might show more sensitivity to changes in terms of response speed. In terms of response intensity, they might show relatively attenuated neural response, due to the nonlinear relation between theta/beta ratio and HRV.

Summarizing, sensory stimuli do not reflect the simple pattern of immediate energy; rather they contain focal, contextual and organic components ^2^. The pooled effect of these components determines and maintains a neural trace of the contextual effect, where the focal component of a deviant stimulus impinges upon organisms already adapted to the context ^2,3,33^. This adaptation is shown to be related to affective experience. Affective levels are established even more easily and quickly through the pooling and interaction of contextual and organic components ^2^. These interactions are reflected in the physiological operations resulting from the functional integration of brain and body ^1^. This view is consistent with Darwin’s proposal that there exists mutual action and reaction between heart and brain, in which emotion is evolved and adaptive ^34^. Although our data did not support the regulation function directly from the heart, we did find that neural response intensity and speed of detecting changes are associated yet independent in the sense that they might be associated with brain-body regulatory coordination of environmental elements. Therefore, assessing perception and adaptation to ambient environmental changes requires a more holistic perspective including context along with biological regulation capacities.

## Competing Interests

**The authors declare no competing financial interests.**

## Supplementary Materials

### Materials and Methods

#### Participants

Twenty right-handed students (70 % females, age *M* = 20.38, *SD* = 2.64) from Cornell University participated in this single session study. Exclusion criteria were any open or healing wounds on the scalp, use of any medication that could affect nervous system processing and any history of neurological disorders. The study was approved by the Cornell University IRB. Informed consent was obtained from each participant and they were compensated with either class credits or $20.

#### Experimental Paradigm and EEG/ECG Acquisition

The experiment involved a passive auditory MMN paradigm that involved listening to auditory stimuli while watching a silent video. In our setup, participants sat comfortably, with limited movement, 75 cm in front of a screen inside an acoustic booth that was insulated from external sounds. They were instructed to ignore the auditory stimuli and instead focus on a silent (with subtitles) BBC documentary of their own choice.

The auditory paradigm, as per a classic MMN paradigm, involved the presentation of 1260 sound stimuli as a sequence of ‘standard’ stimuli (80%) randomly interspersed by ‘deviant’ stimuli (20%). The aim was to compare the neural response to a dynamic change in sound: rising vs. falling pitch. In order to capture this, we designed two sequence conditions: i) ascending MMN sequence, and ii) descending MMN sequence. In these sequences, each standard stimulus consisted of 5 concatenated tones with either rising pitch (600 Hz, 800 Hz, 1000 Hz, 1200 Hz, 1400 Hz) or falling pitch (1400 Hz, 1200 Hz, 1000 Hz, 800 Hz, 600 Hz) respectively. Each tone burst was 50 ms in duration including a rising and falling edge of 10 ms, making each sound stimulus 250 ms. In each sequence, the deviant stimuli were created by replacing the last tone burst by either a higher or a lower pitch than its corresponding standard stimuli i.e. 1600 Hz instead of 1400 Hz for ascending sequence and 400 Hz instead of 600 Hz in descending sequence (illustrated in Fig. 1A). The inter-stimulus interval was fixed at a constant 600 ms. Each type of sequence (ascending and descending) was divided into 7 runs, 2-3 min each. Each run consisted of 144 standards and 36 deviants maintaining the 80/20 ratio. In each run, the first 10 stimuli were specifically standards (to establish the sequence expectation), followed by a pseudo-random distribution of standards and deviants, avoiding consecutive deviants. The runs of the two sequence types were randomly selected with at most two consecutive runs from the same type.

Referential EEG was recorded non-invasively using 128-channel BioSemi active EEG/EMG system (https://www.biosemi.com/products.htm) at a sampling rate of 512 Hz. ECG was recorded from three BioSemi electrodes placed on left and right abdominal region and the region below the right collarbone. Stimuli were presented via BCI2000 software (https://www.bci2000.org/mediawiki/index.php/Main_Page) at 55dBA via speakers placed 40 cm in front of the participant. Precise stimuli onset was recorded by a TTL pulse delivered by an embedded system designed on a 32ARM Cortex-M4 72 MHz CPU that responds in real-time to a sound stimulus above a pre-defined threshold.

#### Data Analysis

EEG was re-referenced to the algebraic average of left and right mastoids, a notch filter (55-65 Hz) and a bandpass filter of 0.5-10 Hz was applied to the raw signal. Bad channels were identified and spherically interpolated. Data was epoched into −100 to 350 ms trials, where time 0 was defined as the time of SOA (stimulus onset asynchrony which was 200 ms after the start of the complex tone pattern, which is also the start of the last tone). Those epochs with obvious abnormal signal segments were visually inspected and manually excluded. The artifacts due to eye movements, muscle and cardiac activity were identified and removed with Independent Component Analysis (ICA), using the EEGLab toolbox ^35^

##### MMN evoked response

The epochs were averaged across trials for each channel to obtain the evoked response. The pre-stimulus period was used for baseline correction. The MMN evoked response was defined as the difference of the evoked response to the standard and the deviant stimuli. We used the coefficient-of-determination (r^2^) to quantify the difference between standard and deviant responses in each time point across the epoch, and then individually selected the channel that has the largest negative r^2^ peak among the fronto-central channels within 80-200 ms for every subject.

##### EEG source localization

Current source densities in each voxel between two sequence conditions (ascending vs descending) were compared by randomization tests on paired data (5000 permutations). Based on statistical non-parametric mapping (SnPM; for details see Holmes et al., 1996 ^36^) corrected for multiple comparison, we used the sLORETA software (http://www.uzh.ch/keyinst/loreta.htm) to perform “non-parametric randomization” of the data.

##### The frontal theta/beta ratio

We estimated the theta frequency band (4-7 Hz) power and the beta frequency band (14-30 Hz) power in the frontal region covered by Afz, Fz, and FCz. The auto spectrum of these channels was calculated using the Welch’s method with a Hamming window of 2 s and 50% overlap. Then the ratio of the theta and beta power was separately calculated for those channels before averaged across channels. The frontal theta/beta ratio was calculated for each sequence condition and averaged across runs.

##### Heart rate variability

We also estimated the heart rate variability (HRV) using the root mean square of the successive differences (RMSSD). It was calculated by taking the square root of the mean squared differences of successive peak to peak intervals, which reflects the beat-to-beat variance in heart rate and is the primary index of vagal mediated heart rate variability ^37^. RMSSD was calculated for each participant across all runs under ascending and descending sequence conditions respectively.

##### Statistical analysis

MMN amplitude did not attenuate and MMN latency did not prolong/extend over runs. We averaged across 7 runs in each condition. The paired t-test was applied to examine the MMN amplitude and latency responses between ascending and descending sequences. To investigate the moderation effects of theta/beta ratio and HRV on MMN amplitude and latency, linear mixed effect models were conducted respectively. High and low theta/beta ratio and HRV were plotted ±1 SD from the mean for descriptive purposes only in the Figures. Inferential analyses maintained the continuous nature of the theta/beta ratio variable.

---

**Fig. S1.**
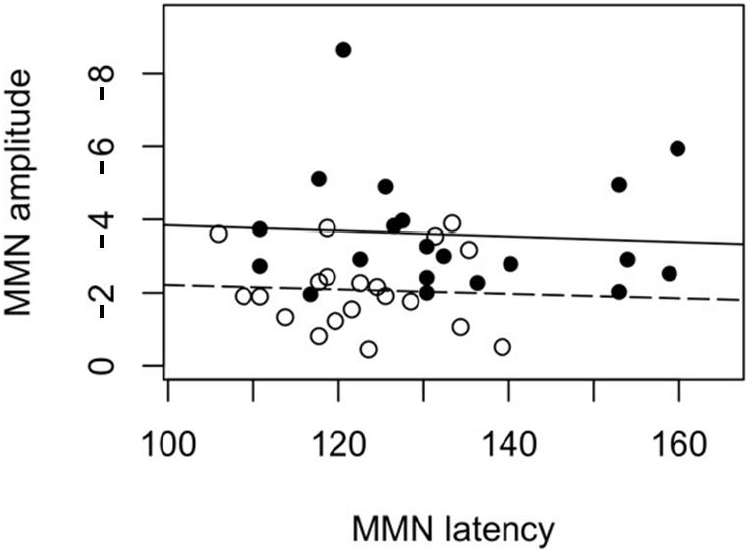
Correlation between MMN latency and amplitude. Across conditions, *r* = 0.14, *t* (38) = 0.83, *p* = 0.4097. In ascending condition, *r* = −0.07 *t* (18) = −0.30, *p* = 0.7664; in descending condition, *r* = −0.05 *t* (18) = −0.22, *p* = 0.8259.

**Fig. S2.**
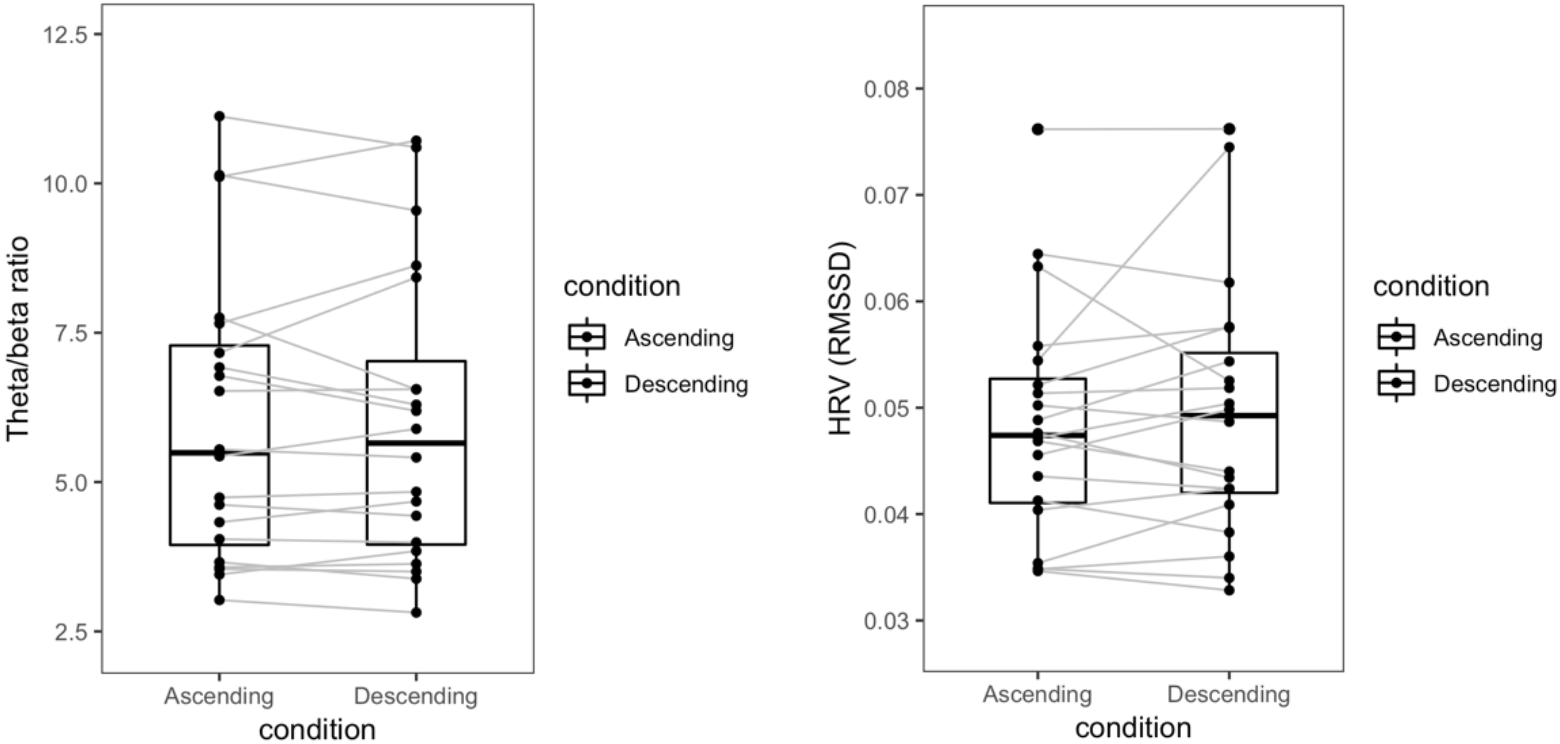
Theta/beta ratio and HRV in two conditions. Paired *t*-test, theta/beta ratio: *t* (19) = −0.77, *p* = 0.4484723, RMSSD: *t* (19) = 0.08, *p* = 0.935995

